# Changes in the gene expression profile during spontaneous migraine attacks

**DOI:** 10.1101/2020.01.08.898239

**Authors:** Lisette J.A. Kogelman, Katrine Falkenberg, Alfonso Buil, Pau Erola, Julie Courraud, Susan Svane Laursen, Tom Michoel, Jes Olesen, Thomas F. Hansen

**Affiliations:** Danish Headache Center, Department of Neurology, Rigshospitalet Glostrup, Glostrup, Denmark; Institute for Biological Psychiatry, Mental Health Center Sct. Hans, Denmark; MRC Integrative Epidemiology Unit, University of Bristol, UK; Department of Clinical Biochemistry and Immunology, Statens Serum Institute Copenhagen, Copenhagen, Denmark; Computational Biology Unit, Department of Informatics, University of Bergen, Norway; Novo Nordic Foundation Centre for protein research, Copenhagen University, Copenhagen, Denmark

## Abstract

**Objective:** Migraine occurs in clearly defined attacks and thus lends itself to investigate changes during and outside attack. Gene expression fluctuates according to environmental and endogenous events and therefore is likely to reveal changes during a migraine attack. We examined the hypothesis that changes in RNA expression during and outside of a spontaneous migraine attack exist which are specific to the migraine attack.

**Methods:** We collected blood samples from 27 migraine patients during an attack, two hours after treatment with subcutaneous sumatriptan, on a headache-free day and after a cold pressor test. All patients were deeply phenotyped, including headache characteristics and treatment effect during the sampling. RNA-Sequencing, genotyping, and steroid profiling was performed on all samples. RNA-Sequences were analyzed at gene level (differential expression analysis) and at network level, and we integrated transcriptomic and genomic data.

**Results:** We found 29 differentially expressed (DE) genes between ‘attack’ and ‘after treatment’, after subtracting non-migraine specific genes, i.e. genes related to a general pain/stress response. DE genes were functioning in fatty acid oxidation, signaling pathways and immune-related pathways. Network analysis revealed molecular mechanisms affected by change in gene interactions during the migraine attack, e.g. ‘ion transmembrane transport’ and ‘response to stress’. Integration of genomic and transcriptomic data revealed pathways related to sumatriptan treatment, i.e. ‘5HT1 type receptor mediated signaling pathway’.

**Interpretation:** Using a paired-sample design, we uniquely investigated intra-individual changes in the gene expression during a migraine attack. We revealed both genes and pathway potentially involved in the pathophysiology of migraine.

## Introduction

The promise of genetic studies, using for instance genome-wide association ^1^ or family studies ^2,3^, to understand the molecular mechanisms of migraine have not been fulfilled and have not led to novel drug targets. An alternative approach is to study the fluctuating gene expression, which is driven by both genetic and environmental factors ^4^. A few gene expression studies indicated a difference between migraine patients outside of migraine attack and healthy controls ^5–7^, but these studies were explorative and after correction for multiple testing, we have recently showed that there is no distinct difference outside of attack ^8^. Only one study has investigated gene expression during migraine attack ^9^. This study compared the migraine patients during attack to healthy controls which is not an optimal design because of huge interindividual gene expression variability. Given this study design it was not possible to establish whether the gene expression was altered due to attack or other clinical characteristics with a reasonable precision although some interesting findings, such as the altered expression of platelet-related genes, supported further study of RNA expression in migraine.

The fluctuating feature of gene expression over time enables a temporal design, i.e. following gene expression changes during fluctuation of disease symptoms. Thus, analyzing individual temporal changes provides the opportunity to study the molecular mechanisms involved in a migraine attack ^4^. However, to retrieve sequential samples during and outside a migraine attack is a demanding task, necessitating a highly organized approach and considerable manpower, and has not previously been reported. The research facilities at the Danish Headache Center enabled such a design. Therefore, we here investigated the intra-individual changes in gene expression during a migraine attack which lasts between four and 72 hours, enabling a paired-sample design. This design increases the statistical power substantially as factors with medium to high variation at an individual level that are not related with the migraine attack are strongly diminished ^10^.

The specific aim of the present study was to investigate gene expression profiles using RNA-sequencing of migraine patients during a migraine attack, two hours after receiving acute medication, and outside migraine attack. We analyze the data on a gene-level as well as on gene network-level and investigate the effect of genomic variations on gene expression alterations. Gene expression changes during a cold pressor test in the same individuals, enabled subtraction of genes putatively involved in general pain/stress response. Our hypothesis was that there are specific changes in RNA expression during migraine attack that are not seen during cold pressor test induced pain.

## Methods

### Study population

We included 17 migraine without aura (MO) and 10 migraine with aura (MA) patients in the study. The MA patients could also have some MO attacks. Inclusion criteria were: diagnosed according to IHS criteria ^11^, female, aged 18-70 years, weighing 45 to 95 kg and of Danish ethnicity. Exclusion criteria were any recent change in daily medication, pregnancy and breastfeeding. All included patients had a full medical history taken at the hospital. Electrocardiography (ECG), physical examination, vital signs and a validated semi-structured headache questionnaire were also conducted at the visit. The patients were recruited via the website “forsøgsperson.dk”, via the Danish Headache Centre, via Facebook and by advertising at hospitals.

### Approvals

The study was approved by the ethics committee of Copenhagen (H-15006298), by the Danish Data Protection Agency (I-suite 03786) and the study is registered on clinicaltrials.gov (NCT02468622). All participants gave written informed consent after receiving oral and written information.

### Study procedure

The patients were instructed to contact the responsible doctor or medical student by phone at the onset of a migraine attack. The patient either went to the hospital by taxi or the doctor/medical student went to the patient’s home. A blood sample was taken from the cubital vein immediately after arrival and the patient was subsequently treated with subcutaneous sumatriptan. One patient chose to take a rizatriptan (10 mg) tablet instead of subcutaneous sumatriptan as acute medication. Another blood sample was taken 2 hours after treatment. To collect attack-specific phenotype data, headache intensity, headache characteristics and associated symptoms were noted down during the 2 hours after receiving treatment with time intervals of 30 minutes. Since gene-expression is affected by sex, we only included female patients in this paper.

Approximately a month later around the same time in the menstrual cycle and at the same time during the day as sampling during the migraine attack, and when the patient was headache-free for at least 24 hours (and migraine free for 5 days), we collected a headache-free sample. The blood sampling on the headache-free day was in the same physical place as the attack-sample. Subsequently, a cold pressor test was performed: the subject kept her hand for as long as tolerated (maximum 10 minutes) in ice water and after 60 minutes another blood sample was taken. The design is visualized in Figure 1.

**Figure 1.**
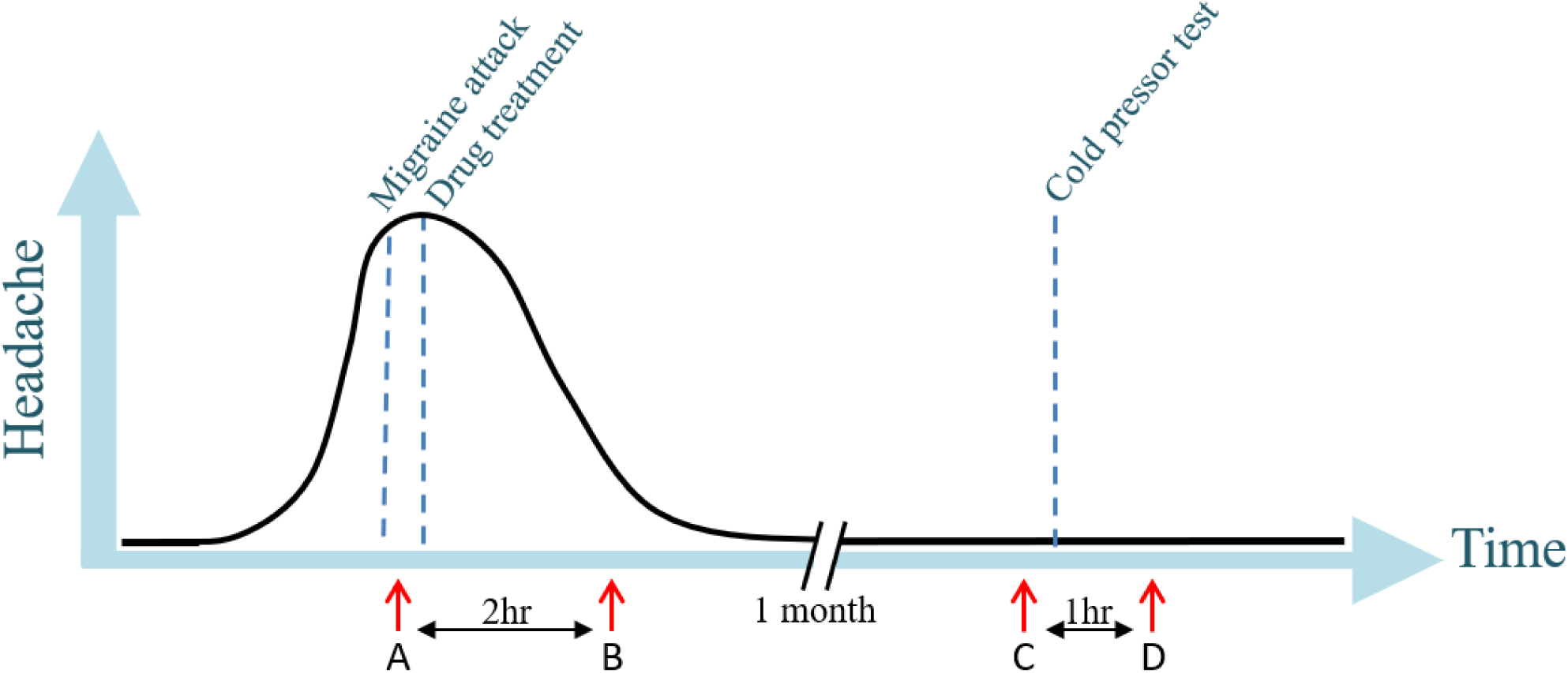
Study design with on the x-axis the time and on the y-axis the degree of headache. Blood sampling (marked by red arrows) was performed during migraine attack (A), two hours after treatment (B), one month later outside attack at a headache-free day (C) and after performing a cold-pressor test (D).

### Steroid level measurement

Solvents were LCMS-grade and purchased from Thermo Fischer Scientific (Waltham, MA, USA) or Sigma Aldrich (St. Louis, MO, USA). Ultrapure water (H_2_O) was obtained on a Milli-Q IQ 7000 LC-Pak (Merck KGaA, Darmstadt, Germany). We used the targeted LCMS CHS MSMS Steroids Kit (PerkinElmer Inc., Waltham, MA, USA) to measure the concentration of 17-hydroxyprogesterone, testosterone, androstenedione and cortisol in the plasma samples. Each experimental sample was extracted once and randomized over the three 96-well plates. We followed the manufacturer’s instructions for sample preparation and data processing. Data were acquired on a Acquity UPLC coupled to a Xevo TQ-S mass spectrometer (Waters Corporation, Waltham, MA, USA). Target Lynx (Waters Corporation) was used to process the targeted LCMS data. Calibration and controls quality met the requirements defined by the manufacturer and by the Clinical and Laboratory Standards Institute (CLSI) guidelines C62-A, paragraph 7.3 (CLSI, 2014). Cortisol (after square root transformation) and androstenedione (after log transformation) were normally distributed, progesterone and testosterone were not normally distributed. In case of normal distribution, a t-test was performed to test for difference between time points. In case of non-normal distribution, a Kruskal-Wallis test was performed.

### RNA-Sequencing and processing

RNA-Sequencing and processing was performed as previously described ^8^. In short, blood samples were stored in PAXgene Blood RNA Tubes and RNA was extracted using the PAXgene Blood RNA Kit (Qiagen, Venlo, The Netherlands) by deCODE Genetics, Reykjavik, Iceland who also performed RNA-Sequencing. Quality of RNA was checked using the Agilent 2100 Bioanalyzer and LabChip. The RNA integrity number (RIN) had a mean of 7.75 (range 6.49 – 9.01). RNA-Sequencing was performed on the Illumina Novaseq, resulting in paired-end reads of 125 basepairs long. Files were processed and quantified with kallisto v0.42.5 using the human reference transcriptome (Gencode Release 28). We successfully collected RNA from 27 migraineurs during attack, 25 migraineurs after treatment, 26 migraineurs outside migraine attack at a headache-free day, and 25 after cold-pressor test. Next, transcript abundances were merged into gene abundances using the R package tximport. Normalization was performed using DESeq2, where the matrix was normalized for library size and gene-length bias using the average gene length-matrix resulting from kallisto. Outliers were detected using the Mahalanobis’ distance ^12^ (MD) using the first eight principal components; a MD larger than the chi-square value for df = 8 at an alpha value of 0.01 were removed. Two samples were removed from analysis: one sample two hours after treatment, and one sample after cold-pressor test. Data was normalized using the DESeq2 package ^13^ using the gene-length/sequencing-depth matrix estimated by kallisto. Genes that were not expressed in at least 90% of the samples were removed and only protein-coding genes were retained for analysis (n = 15,940).

### Differential gene expression analysis

Differential gene expression analysis was performed as a paired sample design, extracting samples of two time points at a time. Differentially expressed (DE) genes between the two time points were detected using a binomial Wald Test within the R package DESeq2 ^13^, with individual patient ID as covariate. P-values were adjusted for 15,940 tests using Bonferroni correction (P_adj_) and were called DE in case P_adj_ < 0.05. First, we compared ‘during attack’ with samples taken ‘headache-free’ (i.e. A vs C [Figure 1]). Second, minimizing the time taken between investigated samples, we compared ‘during attack’ with ‘after treatment’ (i.e. A vs B [Figure 1]). To exclude genes that were not migraine-specific but in general pain/stress related we compared ‘headache-free’ with ‘after cold pressor test’ (i.e. C vs D [Figure 1]). To exploit the temporal design, we performed a likelihood ratio test with the full model including ‘Time’ as covariate using the samples taken during attack (A), after treatment (B), and headache-free (C), and the reduced model omitted ‘Time’ as covariate. Again, p-values were adjusted using Bonferroni correction and genes were called DE when Padj < 0.05.

### Gene regulatory network construction

A differential gene regulatory network was constructed using Lemon-Tree version 3.0.5. ^14,15^. As non-varying genes contain limited information in a network, several thresholds were applied to limit the number of input genes: the top varying genes (n = 1,000, 2,500, 5,000, 7,500 and 10,000) were selected based on their coefficient of variation using the four time points. Data was converted to z-scores, scaling to a mean of zero and standard deviation of 1. The different datasets were clustered in 2-100 clusters using k-means clustering. An elbow plot was constructed to examine the percentage of variance explained by the clusters and, based on this plot, network construction was conducted using the 7,500 most varying genes clustered in 20 clusters, capturing ∼33% of the variance, which was at the inflection point of the elbow plot. The ‘revamp’ task of Lemon-Tree was applied to the subsets of the scaled gene expression matrix based on the four time points ^16^. Genes were reassigned to clusters to maximize the Bayesian co-expression clustering score sampling a parametric threshold (threshold = 4 × 10^−4^). With a low threshold many of the genes will be moved from one cluster to another that maximizes the clustering score, while with a high threshold the genes will remain in the cluster. Plotting the overlap of genes in clusters in the two subnetworks (during attack vs after treatment and headache-free vs after cold-pressor test) resulted in a sigmoid curve where we then picked the switching point to ensure a balance in optimization of the clustering score. Next, gene ontology (GO) enrichment analysis was performed on each cluster in the four subnetworks using the ‘go_annotation’ task of Lemon-Tree that utilizes the BiNGO library ^17^. In the preliminary analysis, the absence of significant DE genes in ‘during attack’ vs ‘headache-free’ indicated that we did not have enough power to detect changes with this sample size over a longer time frame. Then, to avoid potential false positive findings, we did not compare ‘during attack’ vs ‘headache-free’ using the network analysis, and we only investigated changes in the network ‘during attack’ vs ‘after treatment’ and subtracted changes in the network ‘headache-free’ vs ‘after cold pressor test’. Migraine attack enrichments were defined as the difference in significant GO terms between the ‘during attack’ and ‘after treatment’ networks, where the adjusted p-value had to be below 0.05 in one of the networks and above 0.10 in the other network, to ensure that differences in GO enrichment were not driven by noise (e.g. a single gene difference). The workflow is visualized in Figure 2. The same was applied to the ‘headache-free’ and ‘after cold-pressor test’ network, to exclude enrichments that were associated with a general pain/stress response.

**Figure 2.**
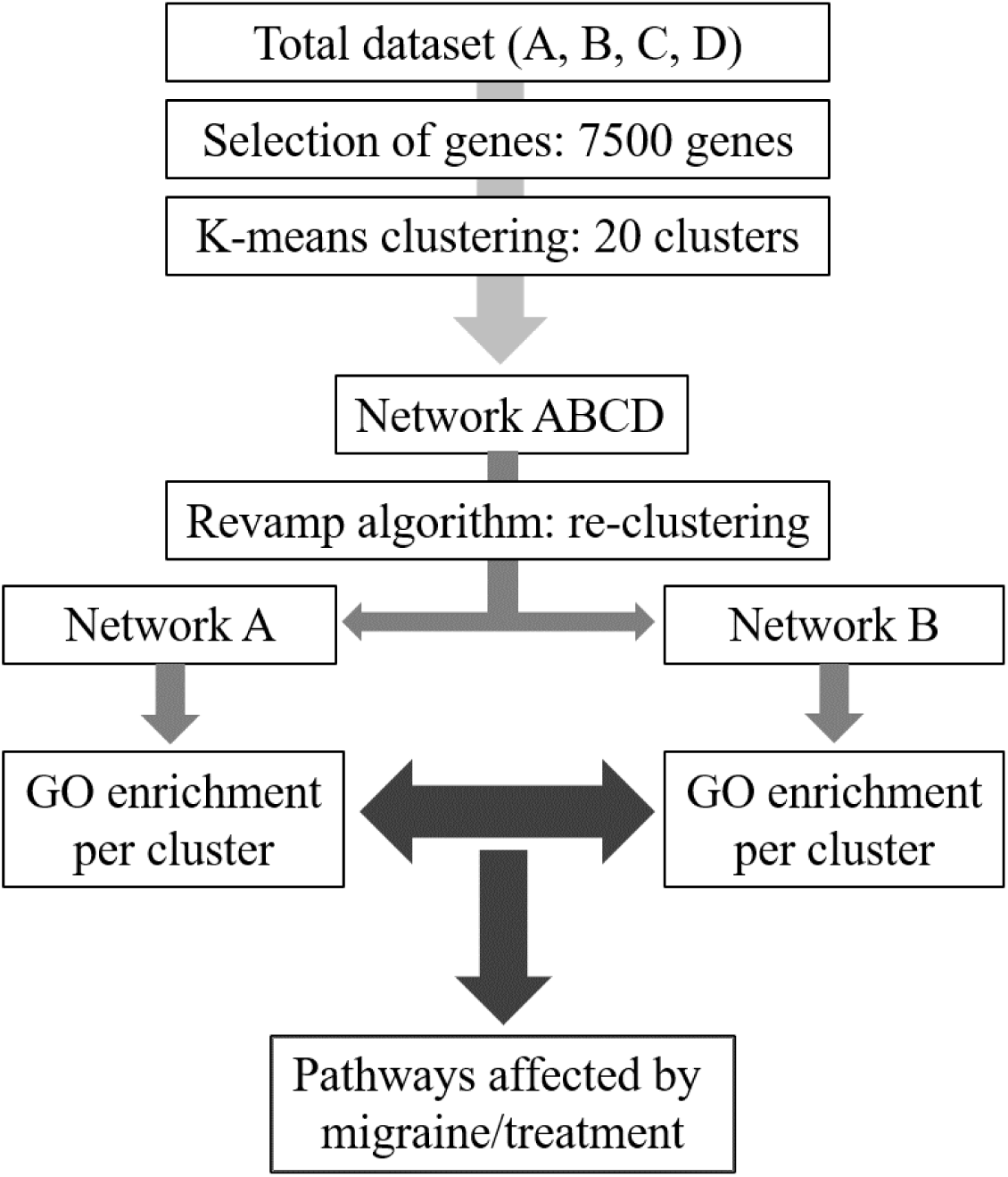
Workflow of differential gene regulatory network analysis, where the different time points are represented by A = during attack, B = after treatment, C = headache-free, D = after cold-pressor test.

### Expression quantitative trait loci (eqtl)

Gene expression changes are being controlled by both environmental factors as well as genetic factors. To investigate which genetic factors (variants) affect gene expression changes during the migraine attack, we integrated genotype data with the transcriptomic profiles. Genotyping was performed using the Illumina HumanOmniExpress 24v1 chip and quality control was performed as described previously ^18^. First, we calculated the change in gene expression by deducting the gene expression profile ‘after treatment’ from the gene expression profile ‘during attack’, both in TPM and in log(TPM). A log transformation was applied to test proportional changes instead of additional changes. Secondly, we ran QTLtools ^19^ using a linear model, investigating all cis interactions using a window of 20Kb and only including SNPs with a MAF > 0.20. The intra-individual variation in drug efficacy may affect gene expression, and consequently detection of eQTLs, and the reduction in headache score (%) was therefore included as covariate. We applied the permutation procedure (n_permutation_ = 1,000) to generate adjusted p-values given by the fitted beta distribution. Adjusted p-values were subsequently corrected using Storey & Tibshirani false discovery rate procedure ^20^ (R/qvalue package).

## Results

### Clinical descriptive statistics

We performed RNA-Sequencing on 27 female migraineurs (17 without aura [MO] and 10 with aura [MA]) longitudinally at four time points: two to measure gene expression during migraine attack (A and B [Figure 1]) and, separated by a month two to measure a general pain-stress response after a cold-pressor test (C and D [Figure 1]). No significant difference was present in attack symptoms (character, aggravation by activity, nausea, vomiting, photophobia, phonophobia) or other characteristics (heart rate, blood pressure) between MO and MA. Time between onset of attack until sample was drawn was lower in MA patients than in MO patients (100 [SD = 47] vs 206 [SD = 98] minutes, respectively; p-value = 1.2 × 10^−3^). Three of the 27 migraineurs (two MO, one MA) did not respond to triptan sufficiently (<50 % reduction of headache after two hours), corresponding to an 89% response rate to triptan (Figure 3). Other changes in migraine characteristics during the migraine attack and after treatment are presented in Table 1. No difference in hormone levels (17-hydroxyprogesterone, testosterone, androstenedione) were present during attack (A vs B [Figure 1]), or between the samples during attack and samples when patients were headache-free (A vs C [Figure 1]).

**Table 1.**
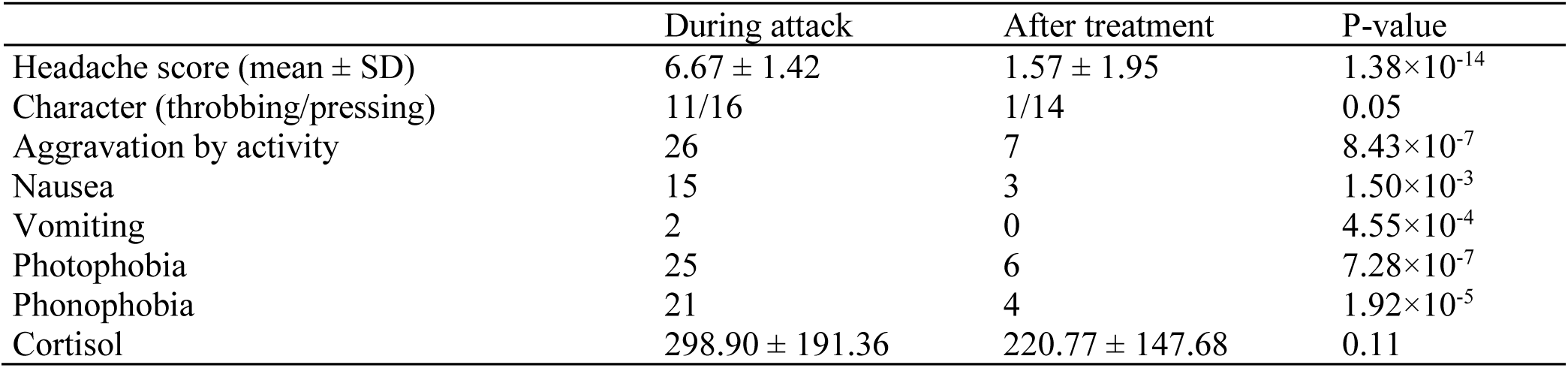
Characteristics during attack and two hours after treatment.

|  | During attack | After treatment | P-value |
| --- | --- | --- | --- |
| Headache score (mean ± SD) | 6.67 ± 1.42 | 1.57 ± 1.95 | 1.38$\times 10^{-14}$ |
| Character (throbbing/pressing) | 11/16 | 1/14 | 0.05 |
| Aggravation by activity | 26 | 7 | 8.43$\times 10^{-7}$ |
| Nausea | 15 | 3 | 1.50$\times 10^{-3}$ |
| Vomiting | 2 | 0 | 4.55$\times 10^{-4}$ |
| Photophobia | 25 | 6 | 7.28$\times 10^{-7}$ |
| Phonophobia | 21 | 4 | 1.92$\times 10^{-5}$ |
| Cortisol | 298.90 ± 191.36 | 220.77 ± 147.68 | 0.11 |

**Figure 3.**
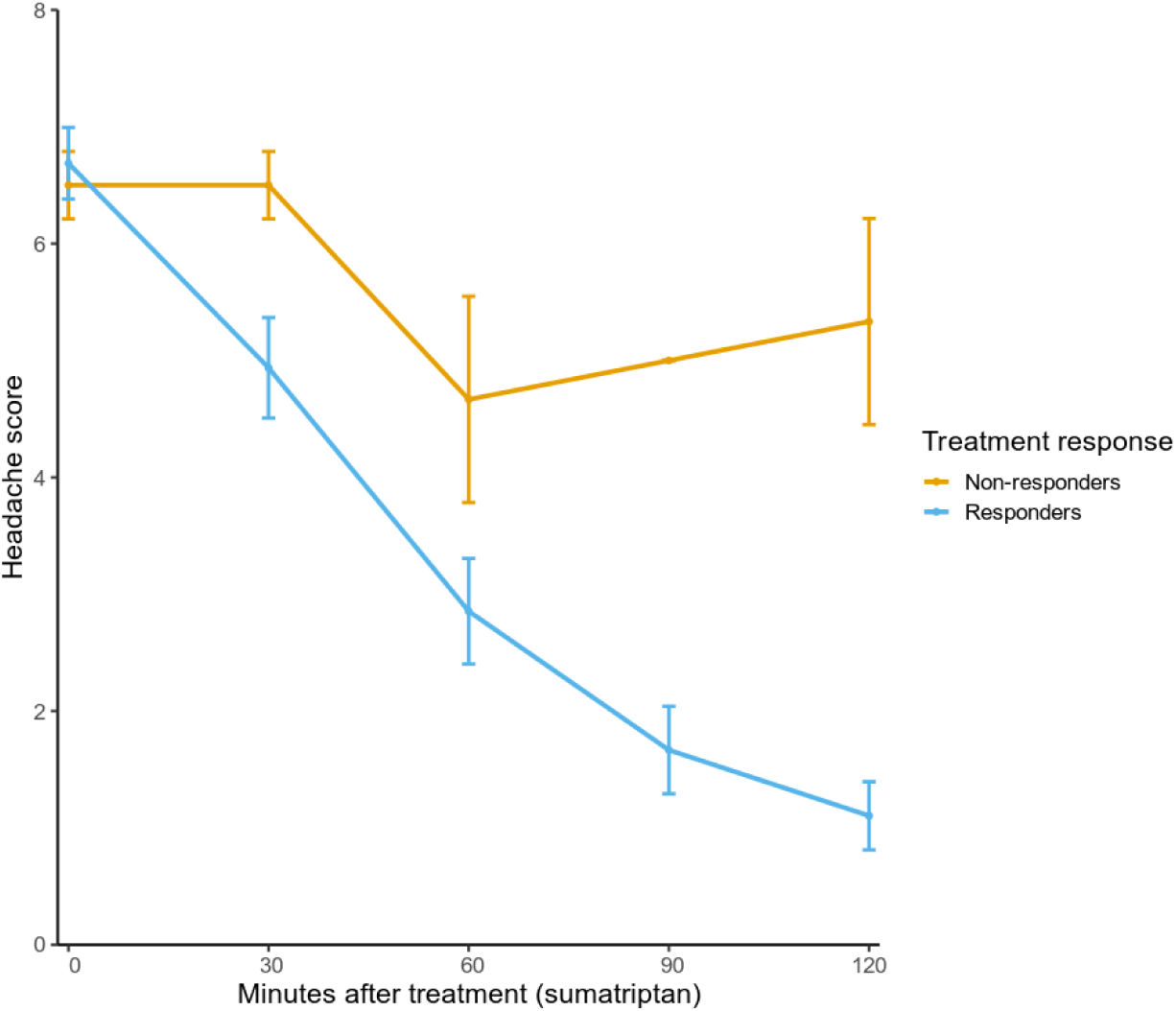
Difference in headache during the two hours after treatment with sumatriptan.

### Differential expression

Gene expression analysis of 15,940 genes comparing ‘during attack’ versus ‘headache-free’ (A vs C [Figure 1]) did not identify any significantly differentially expressed (DE) genes after multiple-testing correction, with the top genes being *TNFRSF17* (P = 6.50×10^−5^), *NOSTRIN* (P = 1.76×10^−4^) and *LRRTM2* (P = 3.29×10^−4^). We identified 33 significantly DE genes (P_adj_ < 0.05) between ‘during attack’ and ‘after treatment’, i.e. A vs B (Figure 1). All DE genes are presented in Table 2. Of those, twenty were down-regulated after treatment and thirteen were up-regulated. To incorporate the sample taken ‘headache-free’, we fitted a temporal model, which confirmed four of the DE genes (*CPT1A, ETFDH, SLC25A20*, and *SLC46A2*), see Figure 4.

**Table 2.**
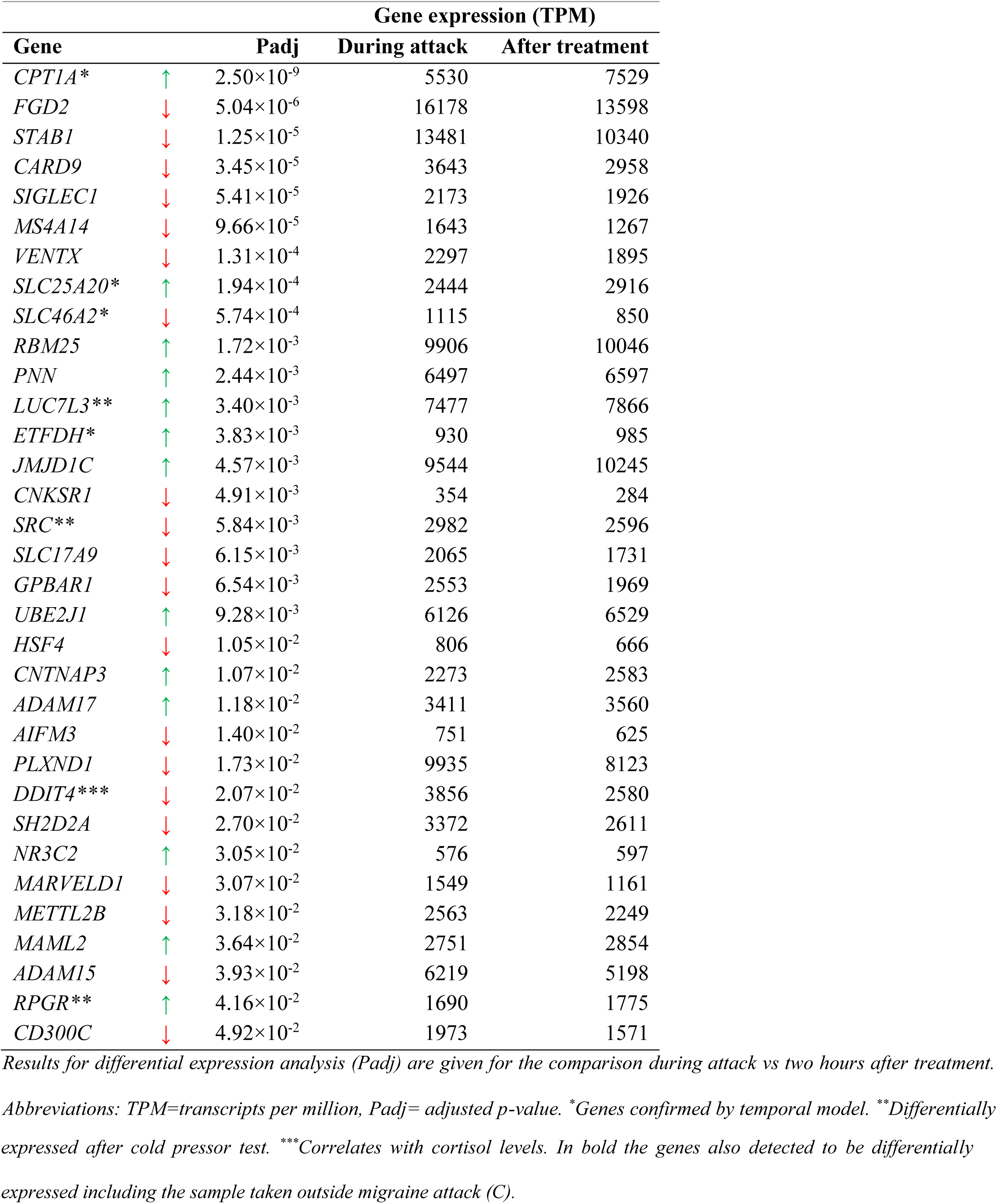
Genes differentially expressed during a migraine attack.

**Figure 4.**
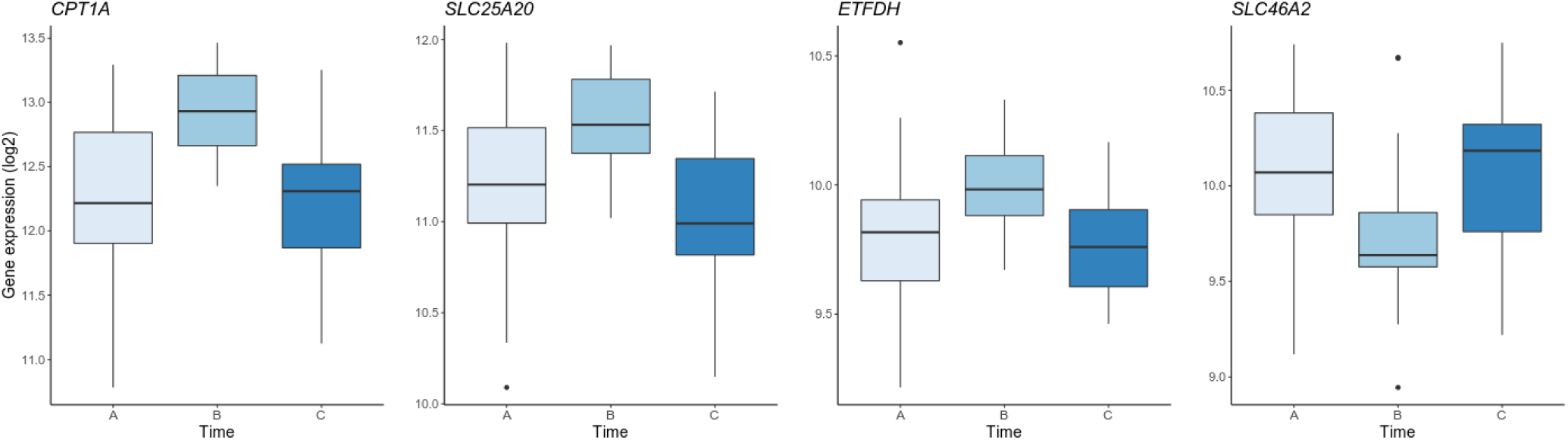
Expression of the four differentially expressed genes, detected both in the paired-sample design (i.e. A vs B) and in the temporal model (i.e. with timepoint of sampling as covariate). On the y-axis the gene expression in transcripts per million (log2) and on the x-axis the three time points (i.e. A = during attack, B = after treatment, C = headache-free).

Genes that are altered during the cold pressor test are expected to be involved in a general pain/stress reaction and are not migraine-specific. Three of the DE genes were DE after cold pressor test (i.e. C vs D [Figure 1]) and were therefore not considered to be migraine genes (*LUC7L3, SRC* and *RPGR*). Although the cortisol levels did not significantly decrease after treatment, we did see one of the DE genes to be significantly correlated with cortisol levels: *DDIT4* (r = 0.78, P_adj_ = 9.11×10^−10^). *DDIT4* is known to be induced by cortisol ^21^ and known to be expressed under stress ^22^. None of the DE genes significantly correlated with one of the other steroids measured.

The migraine patients were treated with subcutaneous sumatriptan and thus we tested whether the DE genes were enriched for genes that could be affected due to the treatment. Triptans are serotonin agonists ^23^, and therefore, we investigated genes within the serotonergic synapse (KEGG pathway hsa04726). We did not see an overall differential expression within the pathway, (P = 0.62), but one gene was significantly DE: *BRAF* (P = 3.79×10^−4^). As seen from Figure 2, there were patients that did not respond sufficiently to treatment and this may have affected the differential expression analysis. Excluding those patients (headache reduction < 50%) does reduce the statistical power, but all DE genes were still DE and, consequently, the DE analysis does not seem to be affected by non-responders.

The most recent genome-wide association study (GWAS) has shown 38 loci pinpointing to 51 genes involved in the susceptibility to migraine ^24^. We tested whether those genes were altered during migraine attack: two genes were significantly DE during migraine attack (A vs B [Figure 1]) and not after cold pressor test (i.e. C vs D [Figure 1]): *LRP1* (P = 1.30×10^−5^) and *TSPAN2* (P = 2.16×10^−4^). None of the genes were DE when including the headache-free sample in the analysis using the likelihood test. Expression of GWAS genes during migraine attack are presented in Supplementary Table 1.

### Gene regulatory network

We constructed a differential gene regulatory network, enabling us to assess changes in the interaction between genes. If a change in gene interaction during migraine attack (from network ‘during attack’ [A] to network ‘after treatment’ [B]) led to a significant change in pathway composition, based on gene ontology (GO) enrichment, we considered the pathways affected by, or interacting with, the migraine attack or its treatment. Changes in pathway composition after cold pressor test were excluded to omit general pain/stress response related changes. We found 270 migraine attack related GO terms (Supplementary Table 2), which were enriched during attack (A) or after treatment (B). Among the most significant differential GO terms enriched during attack (A) is ‘ion transmembrane transport’ (cluster 13) and among the most significant GO terms enriched after treatment (B) is ‘response to stress’ (cluster 2). Cluster 14 was enriched of DE genes (n=8, P = 2.84×10^−5^), with enrichment of immune-related GO terms (e.g. antigen processing and presentation via MHC class Ib, B cell mediated immunity), hemopoiesis and gluthathione derivative metabolic process. Of the 41 genes targeted by risk variants from the migraine GWAS, 24 were present in the differential network, for example *ECM1* and *MPPED2* in cluster 9, with GO-enrichment of ‘detection of stimulus involved in sensory perception’ during attack (A).

### Expression quantitative trait loci

Integration of genotype with the transcriptomic changes during migraine attack did not reveal any significant eQTLs (qvalue < 0.05). The eQTL analysis undermined the paired-sample design, and consequently the small sample size limits the power to detect eQTLs. Still, we applied enrichment analysis over the top eQTLs. Several pathways relevant to migraine were detected: ‘5HT1 type receptor mediated signaling pathway’, ‘metabotropic glutamate receptor group II pathway’, ‘dopamine receptor mediated signaling pathway’, ‘muscarinic acetylcholine receptor 2 and 4 signaling pathway’ and ‘histamine H2 receptor mediated signaling pathway’ were nominally significantly enriched (P < 0.05) among the 181 top eQTLs (P_beta_ < 0.01). Based on proportional changes, the top 161 eQTLs (P_beta_ < 0.01) were nominally enriched in the ‘cholesterol biosynthesis’ pathway (P = 1.18 × 10^−2^).

## Discussion

Though genetic factors play a major role in migraine pathophysiology, the involved genetic mechanisms are largely unknown. Using RNA-Sequencing we investigated the changes in gene expression during a migraine attack and revealed genes and pathways that are potentially involved in the underlying mechanisms of migraine. Thirty-three differentially expressed genes were found and, using a network approach, several potential migraine mechanisms were pinpointed. Furthermore, by integration of genomic and transcriptomic data we showed changes due to pharmacological treatment.

### Changes at gene level during the migraine attack

Among the up- and down-regulated genes during migraine attack were genes functioning in fatty acid oxidation (*CPT1A, SLC25A20* and *ETFDH*), signaling pathways such as Notch signaling (*MAML2, ADAM15* and *ADAM17*) and immune-related pathways (*CARD9, SH2D2A, CD300C*). Fatty acid oxidation is the process of breaking down a fatty acid into acetyl-CoA units, taking place in mitochondria. It has been implicated to play a role in migraine pathophysiology. For example, one of the migraine-GWAS hits is linked to *ACSL5*, which activates long-chain fatty acids ^25^. Also, mitochondrial dysfunction has been suggested in migraine ^26,27^. Those mechanisms are also translated into migraine treatment: dietary intake of fatty acids was associated with a lower frequency of migraine attacks ^28,29^, and riboflavin, which affects the mitochondrial energy efficiency, is used as a prophylactic treatment ^30,31^. Also signaling pathways were present among the DE genes. One of the Notch signaling genes, *MAML2*, was also differentially expressed in cluster headache patients ^32^. The Notch signaling pathway has previously been associated with migraine, among others due to the GWAS hit near *NOTCH4* ^1^ and a mutation in *NOTCH3* results in CADASIL (cerebral autosomal dominant arteriopathy with subcortical infarcts and leukoencephalopathy) with migraine as one of the main symptoms. Furthermore, several immune genes were up- or down-regulated during the migraine attack, though previous studies have not been able to give straightforward evidence of a relation between the immune system and migraine ^33^.

Four genes were confirmed in the temporal model: *CPT1A, SLC25A202, ETFDH*, and *SLC46A2*. Remarkedly, the first three are all functioning in fatty acid oxidation; the latter encodes a membrane transport protein with unknown function. Furthermore, the eQTL results pointed to cholesterol biosynthesis; as the acetyl-CoA used for cholesterol biosynthesis is derived from fatty acid oxidation this suggests the involvement of fatty acid oxidation during a migraine attack.

### Changes in gene interactions during the migraine attack

Analysis of the system as a whole, using a network approach, gave different insights in processes involved during the migraine attack. Many gene ontology terms were overrepresented in one of the clusters covering a wide variety of processes related to the migraine attack and/or response to migraine treatment. For example, one of the most significant terms active during attack was ‘ion transmembrane transport’. Causal mutations in genes resulting in Familiar Hemiplegic Migraine, a mendelian subtype of migraine, are all involved in ion transport ^34^. Also in common migraine, ion channels are expected to play an important role ^35,36^. Similar to the DE analysis results, several GO terms were immune-related again implying a role of the immune system during the migraine attack. Even though network approaches are dependent on parametric thresholds, we showed that the software Lemon-Tree was able to pick up changes in the network-structure of the gene expression profiles due to biological processes, which both supported and complemented findings from the more-traditional differential expression analysis.

### Treatment of attacks with sumatriptan affects gene expression profiles

Samples were collected during a spontaneous migraine attack and patients were treated immediately after blood sampling with subcutaneous sumatriptan, a very efficient and fast-working migraine treatment. Therefore, the genes and pathways proposed by this study may be derived from a pharmacological effect that is not related to migraine *per se*. Triptans are known 5-HT_1B/D_ (serotonin) receptor agonists, but the exact mechanisms of action are unknown. Our results showed that expression levels of genes known to be involved in the serotonergic synapse were altered during attack (i.e. *BRAF* was DE in our study). Further, the ‘5HT1 type receptor mediated signaling pathway’ was overrepresented among the top eQTLs. A future cross-over design with both a placebo and triptan treatment or treatment outside migraine attack would be able to distinguish between pharmacological and migraine-specific mechanisms.

### Previous findings: expression of platelet genes

Migraineurs during attack (n = 7) were previously compared to healthy controls; 40 upregulated genes were detected in the migraineurs, and it was suggested that platelet abnormalities play a role in migraine ^9^. In our study, the genes present in the KEGG pathway ‘platelet activation’ were not significantly DE (P = 0.18), neither were the DE genes platelet-related. However, of the 40 upregulated genes, 31 were expressed in our sample and 15 were DE (P < 0.05), namely *TRDMT1, CDKN1C, ZFP36L1, CCL5, SPARC, PF4, DUSP1, ITGA2B, CLU, GP1BB, CD247, BRD3, SNRPD3, AKR1B1*, and *PTGS2*. KEGG pathway analysis of those 15 genes resulted in overrepresentation of ‘blood coagulation’ (FDR = 4.93 × 10^−2^). Even though our results are not in uniform agreement, platelet genes may be involved during migraine.

### Strengths and limitations

Changes in the gene expression profile are tissue-specific and highly affected by environmental factors. Both the genetic risk loci found in the most recent genome-wide association study ^1^ and the novel CGRP antibodies showing therapeutic effect on migraine attacks ^37^, suggest that vascular and smooth muscle are involved in migraine pathophysiology. Further, it was only possible to collect peripheral blood, which has also shown to correlate with several nervous system tissues ^38,39^. Consequently, we expect that genes identified in peripheral blood explain part of the pathophysiology of migraine.

Even though we optimized the power using a temporal design focusing on intra-individual changes, we were not able to detect DE genes between the samples taken ‘during attack’ and those taken when the same individual was ‘headache-free’. This may indicate that larger sample sizes are needed to obtain power to detect (smaller) changes. It may also indicate that changes due to migraine were not measurable at this time-scale and that we did detect changes due to treatment. Furthermore detecting eQTLs requires a large sample size (i.e. with a MAF of 0.20 a sample size of 135 is required to have a power of 0.8 for a genome-wide eQTL scan ^40^). However, sampling many migraine patients during an attack is a highly demanding task, which requires substantial manpower, and has not been achieved before. Nevertheless, by sampling during attack and after treatment we minimized intra-individual variation and showed changes in gene expression and pathways. This provided clear hints to the molecular mechanisms during the migraine attack. The subtraction of changes during cold-pressor test ensured the specificity of migraine-relevant genes and pathways over general pain/stress-related genes.

Furthermore, even though underpowered, our genome-wide eQTL scan pointed out the involvement of the serotonergic pathway. As the migraine patients were treated with triptan which acts on the serotonergic pathway, this ensured that we were able to reveal relevant molecular mechanisms using this study design. This encourages coordinated international multi-centers. Future studies may exploit the temporal design further by including multiple samples during the migraine attack and/or sampling more individuals to increase power to e.g. find eQTLs.

In conclusion, our study for the first time investigated intra-individual changes in gene expression profiles using state-of-the-art RNA-Sequencing technology and is therefore methodologically an important step forward in elucidating migraine mechanisms. We showed that the paired-sample design gives the power to reveal molecular mechanisms involved in a migraine attack and its treatment and that the cold pressor test, importantly, excluded changes due to a general stress/pain response. Our study suggests several molecular mechanisms involved in migraine pathophysiology, with an important role of mitochondria, Notch signaling, ion channels, the immune system and previous findings suggesting platelet abnormalities.

## Acknowledgements

We thank all migraineurs for participating to this study. We thank Isabel Engel, Lau U. Poulsen and Jacob Worm for their assistance in collection of samples. We thank Gísli H. Halldórsson, Hreinn Stefánsson and Kari Stefánsson (deCODE genetics, Iceland) for RNA-Sequencing and genotyping of the samples. We thank Madeleine Ernst and Arieh Cohen (Statens Serum Institute) for obtaining steroid levels of the samples. The work was funded by a grant from the Candys foundation ‘CEHEAD’ and EU-funded FP7 “EUROHEADPAIN” grant (no. 602633), both awarded to prof. Jes Olesen. PE was supported by CRUK [C18281/A19169].

## Authors’ contributions

KF and JO conceived and designed the experiments; KF collected the samples. JC and SSL measured steroid levels in the samples. PE and TM assisted with gene regulatory network construction. AB assisted with the integration of genomic and transcriptomic data. LJAK and KF wrote the first draft of the manuscript. All authors read and approved the final manuscript.

